# Evaluating in-context learning with prompting regimes for transformer-based synthetic health tabular data generation

**DOI:** 10.64898/2026.09.29.754354

**Authors:** Amanda Bertgren, Fredrik Öhberg, Paolo Soda, Ulf Näslund, Patrik Wennberg, Christer Grönlund

## Abstract

Tabular synthetic data generation (SDG) can facilitate research and development in the health domain, as access to health data is restricted. To accommodate for its structural complexity, such as multimodal distributions and fragmentation, transformer models provides an alternative to conventional SDG methods since they are better equipped to infer global spatial coherence due to textual context of tabular records. However, methods for optimising their adaption to tabular SDG are still largely unexplored. We contribute to the understanding of transformer-based SDG by evaluating fidelity and privacy of three prompting regimes, zero shot, one shot and few shot training, for tabular health SDG using small transformer models. We evaluated llama 3.2 (1B and 3B parameters) and Phi (1.3B and 2.7B parameters), on a real clinical dataset. Our results showed that zero shot learning was less prone to hallucinations and performed overall better for fidelity, but at the loss of privacy in comparison to the other regimes. Our results showed no differences between model sizes and families. Future studies should extend the analysis to other models and datasets, and explore strategies to reduce hallucinations.

## 1 Introduction

The rise of electronic health records has introduced big data into the healthcare domain [1]. The vast collection of health data can facilitate research and development in different areas of healthcare, such as clinical research or predictive analysis of disease occurrence. Even health digital twins may be possible in the future [2] as the utilisation of health datasets increase.

To enable advances in these areas, challenges linked to the use of health data for research and development purposes must be addressed. One challenge concerns access to the data. Since health data is highly sensitive, regulations such as the General Data Protection Regulation (GDPR) in Europe [3] and the Health Insurance Portability and Accountability Act (HIPAA) in the United States [4] limit access to protect the privacy of individuals. To comply, health data institutes often remove direct identifiers and do a case-by-case assessment before distributing data, but this is often time-consuming and privacy-risk prone [5, 6]. Synthetic data generation (SDG) can be used as an alternative, as synthetic data can be generated with similar statistical properties as the real dataset and the level of privacy can be controlled [7].

Another challenge concerns the low data quality of health data, since it often contains imbalanced labels, fragmented domain data and high proportions of missing data. These conditions limit SDG since SDG methods often utilise generative adversarial networks or autoencoders [7, 8] with an inherent structural inflexibility due to lack of local spatial coherence in tabular data. Recent advances have shown that transformer models with improved techniques for reasoning and trained on large numerical training corpora can constitute an alternative, since these can process global spatial coherence with textual context [9, 10, 11]. However, since transformer models are specialised on textual input and output, optimised formats for handling of tabular data in transformer-based SDG remain largely unexplored.

### 1.1 Contribution

We extend the understanding of transformer-model based tabular SDG by investigating the impact of in-context learning. We do this by evaluating the effect of three prompting regimes, zero shot, one shot and few shot learning for SDG on small (≤ 3B parameters) transformer models on a real clinical dataset. Our results contribute to the foundational understanding of transformer models as we evaluate a pipeline component independently to isolate its effect.

## 2 Data and Methods

### 2.1 Dataset

The dataset was retrieved from the VIPVIZA (VIsualiZAtion of asymptomatic Atherosclerotic disease) study - a randomised controlled trial for individuals at ages 40, 50 and 60 in Västerbotten county, Sweden [12, 13], investigating the effect of pictorial risk communication on risk of cardiovascular disease.

The selected data included *n* = 2603 samples and *m* = 32 variables spanning multiple health domains from the VIPVIZA study, including clinical risk markers, psychology, ultrasound imaging, and lifestyle. The selection of variables was based on the combined recommendation of VIPVIZA experts within the domains. The dataset *D* was sequentially split into train (80%), validation (10%), and test (10%) datasets, *D* = *D*_train_ ∪ *D*_val_ ∪ D_test_, to avoid information leakage.

D was encoded by setting descriptive column names, exchanging coded categorical values with descriptive labels, row-wise delimetered concatenation of columns and wrapping each row with a start and end tag, <row> and </row>.

### 2.2 Transformer models and experiments

We evaluated two model families. We used two models from the llama 3.2 family (sizes 1B and 3B parameters) as well as phi-1.5 (1.3B parameters) and phi-2 (2.7B parameters)) from the phi model family [14, 15].

Three experiments with different prompting regimes were conducted for each model.

1. *Zero-shot prompting*. The prompt was constructed only consisting of an open ended row tag, <row>, to initiate completion by the transformer model.
2. *One-shot prompting*. The prompt was constructed by randomising one sample from _train_ and adding an open ended row tag, <row>, at the end to initiate completion by the transformer model.
3. *Few-shot prompting*. The prompt was constructed by randomising three sample from D_train_, joining them to a string, and adding an open ended row tag, <row>, at the end to initiate completion by the transformer model.

Five datasets were generated per experiment and model to account for stochastic variability, and the evaluation was performed on all five dataset before averaging the results.

### 2.3 Synthetic Data Generation

All models were trained using a per-device batch size of 1 with gradient accumulation over 4 steps. A warmup of 100 steps was applied, and training was run for a maximum of 2000 steps. Mixed precision was enabled using fp16, and the maximum tokenizer length was set to 512 after approximating the number of tokens needed for each row in D_train_ (range from 988 to 1178 characters). We used Low-Rank Adaptation (LoRA) for causal language modelling with rank 16 and scaling factor *α* = 32 on the query and value projection layers. LoRA updates were applied without bias and with a dropout rate of 0.05.

For generation, the temperature was set to 0.8 to increase crativity in small models [16], nucleaus sampling was set to 0.9 to promote decoding diversity and default repetition penalty was set to 1.1 to penalise direct reconstruction.

### 2.4 Evaluation

The models and experiments were evaluated for univariate and bivariate fidelity in numerical and categorical variables. Univariate fidelity was evaluated using the Kolmogorov-Smirnov statistic, *KS*, for numerical variables and Jensen-Shannon divergence, *JS*, for categorical variables. Bivariate fidelity was evaluated using Pearson correlation difference, |*ρ*_*real*_ − *ρ*_*syn*_|, and joint Jensen-Shannon divergence, *JS*_*joint*_, for numerical and categorical variables respectively.

Privacy was evaluated using mean nearest neighbour distance ratio, *NNDR*, median distance to closest record, *DCR* and *ε*-identifiability risk, *ε* − *IR*. All privacy metrics were computed using the Syntheval package [17] and missing values were imputed solely for the computation using −1 for numerical values and the string *“Missing”* for categorical variables due to requirements of the metrics and package.

Any hallucinated columns or labels were excluded in a pre-processing step before evaluation of fidelity and privacy, however, the errors were stored and occurrences were analysed and reported.

## 3 Results

The fidelity metrics across all models and prompting regimes are reported in fig. 1. Overall, zero shot prompting seems to produce better results for the Phi family, whilst few shot prompting seems to perform slightly better for the llama family although less pronounced. For both model families, the size of model did not seem to have an effect on the fidelity results. For univariate fidelity, as reported by 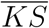 and 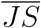, zero shot learning worked better for numerical variables whilst few shot learning produced better results for categorical values. For bivariate fidelity, as reported by |*ρ*_*real*_ −*ρ*_*syn*_| and 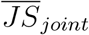, zero shot learning was again better for numerical variables whilst one shot learning displayed better results for categorical variables. However, the standard deviation of numerical bivariate fidelity, | *ρ*_*real*_ −*ρ*_*syn*_|, show that the variability of the results are notably large.

**Figure 1.**
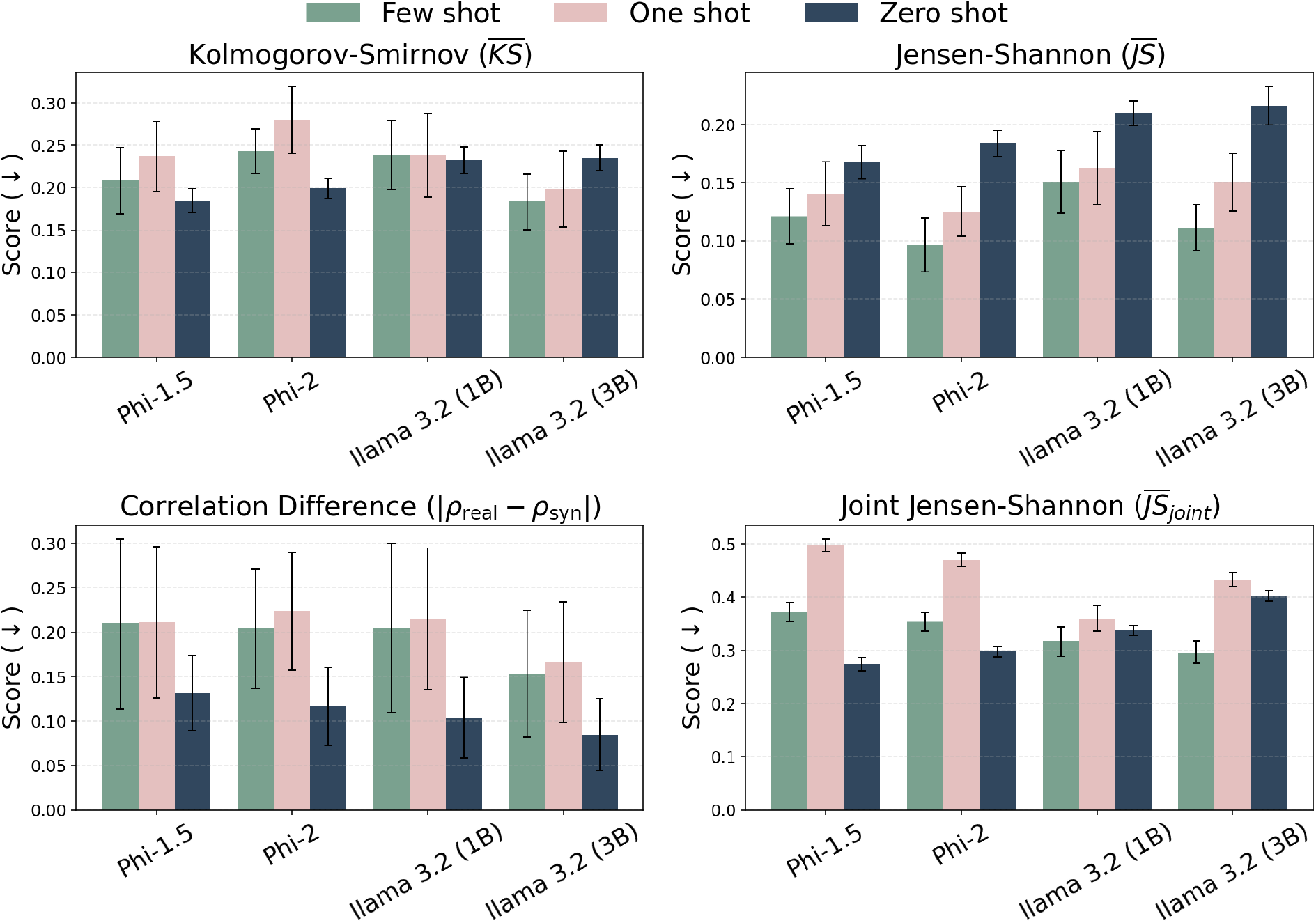
Univariate (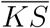 for numerical variables, 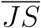 for categorical variables) and bivariate fidelity (|*ρ*_*real*_ − *ρ*_*syn*_| for numerical variables and 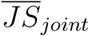 for categorical variables) across all prompting regimes and models.

The privacy is reported in table 1. Overall, zero shot learning performed worse for *NNDR* and *ε* − *IR* than the other prompting regimes, but varied across *DCR*. However, *DCR* generally shows high variability, which may indicate instability in the metric results. Few and one shot learning performed similarly across all privacy metrics, and consistently kept below the recommended threshold of 9% in *ε* − *IR* for all values except for few shot learning with Phi-2, in contrast to zero shot learning which in three out of four experiments exceeded the threshold. The performance across model families and model sizes varied overall and no trend is apparent.

**Table 1:** Privacy metrics across all prompting regimes and models. Values are reported as mean ± standard deviation.

| Model | Shot | $NNDR(\uparrow)$ | $\varepsilon-IR(\downarrow)$ | $DCR(\uparrow)$ |
| --- | --- | --- | --- | --- |
| llama 3.2 (1B) | Few | $0.980 \pm 0.009$ | $0.022 \pm 0.005$ | $0.358 \pm 0.002$ |
| | One | $0.991 \pm 0.004$ | $0.036 \pm 0.004$ | $0.217 \pm 0.198$ |
| | Zero | $0.916 \pm 0.003$ | $0.096 \pm 0.007$ | $1.104 \pm 0.019$ |
| llama 3.2 (3B) | Few | $0.978 \pm 0.004$ | $0.047 \pm 0.006$ | $0.420 \pm 0.064$ |
| | One | $0.940 \pm 0.008$ | $0.015 \pm 0.001$ | $0.565 \pm 0.155$ |
| | Zero | $0.928 \pm 0.004$ | $0.078 \pm 0.006$ | $1.327 \pm 0.021$ |
| Phi-1.5 | Few | $0.987 \pm 0.007$ | $0.011 \pm 0.005$ | $1.353 \pm 0.718$ |
| | One | $0.978 \pm 0.006$ | $0.022 \pm 0.021$ | $0.362 \pm 0.001$ |
| | Zero | $0.902 \pm 0.003$ | $0.106 \pm 0.006$ | $0.987 \pm 0.020$ |
| Phi-2 | Few | $0.966 \pm 0.009$ | $0.139 \pm 0.005$ | $1.356 \pm 0.161$ |
| | One | $0.989 \pm 0.004$ | $0.098 \pm 0.017$ | $2.502 \pm 0.081$ |
| | Zero | $0.905 \pm 0.003$ | $0.114 \pm 0.006$ | $0.937 \pm 0.008$ |

To evaluate the output quality of the models, the averaged error proportion and number of hallucinated columns per model was computed and averaged over the five runs. The error rate was defined as the average proportion of errors per non-hallucinated column for each experiment.

The results are reported in fig 2. Zero shot generation produced nearly no error, except for experiment llama 3.2 (3B). The smaller models showed generally lower error rate, but higher numbers of hallucinated columns. Experiment llama 3.2 (3B) had the overall highest error rate yet overall lowest number of hallucinated columns.

**Figure 2.**
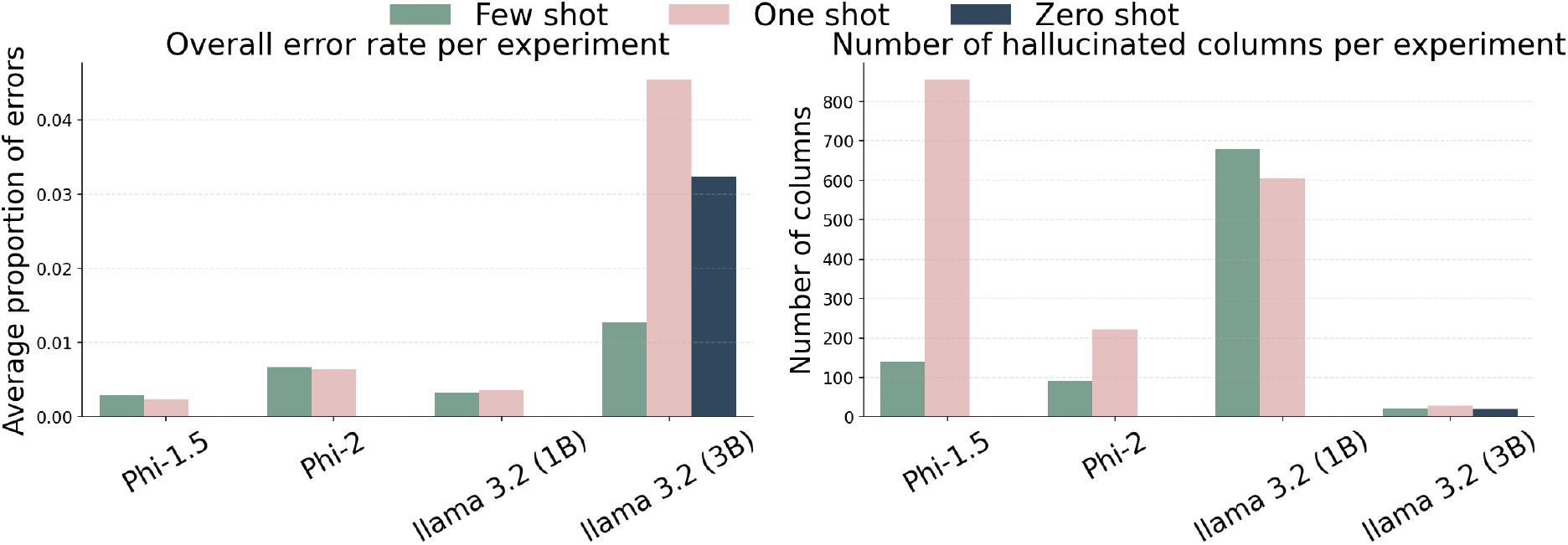
Error rate (averaged over variables and runs) and number of hallucinated columns (averaged over runs) across all prompting regimes and models.

Manual inspection was performed to obtain examples of the errors. The errors consisted of examples such as *“Age 54”*, despite the real age variable only having values in multiples of tens, *“Divorced / Separated / Living alone”* instead of *“Divorced / Separated”* and small typos or changes in lowercase or uppercase letters. The hallucinated columns consisted of examples such as variable names with an additional row tag, like *“<row> Age”*, pure errors, like *“24*.*058399999999971 </block> <block key”* or clinical variables not in the dataset, like*”Drug dosage”*.

## 4 Conclusion

We have evaluated the effect of prompting regimes zero shot, one shot and few shot learning on small transformer models for in-context learning of tabular health SDG. We have shown that the prompting regimes show a trade-off between realism and privacy, similar to the the fidelity-privacy trade off as often observed for SDG. Our results indicate that zero shot learning yields a more robust performance, as seen by lower distributional differences between the real and synthetic data (especially for numerical variables) and fewer occurrences of hallucinations. However, this is at the cost of privacy protection, for which zero shot performed worse than the other prompting regimes and exceeds thresholds for identifiability risk.

Further, our experiments show that the size difference of the models does not seem to affect fidelity or privacy, however, the smaller models seemed to be more prone to hallucination of new columns but less prone to hallucinate new labels. Lastly, the performance was also similar across model families Phi and llama 3.2.

Future studies should build upon this work by extending the analysis to larger transformer models and across more clinical datasets. Strategies for reducing or interpreting hallucinations should also be explored, to avoid removal of near correct values.

